# Occurrence of COVID-19 symptoms during SARS-CoV-2 infection defines waning of humoral immunity

**DOI:** 10.1101/2021.03.26.437123

**Authors:** Jun Wu, Bo-Yun Liang, Yao-Hui Fang, Hua Wang, Xiao-Li Yang, Shu Shen, Liang-Kai Chen, Su-Meng Li, Si-Hong Lu, Tian-Dan Xiang, Jia Liu, Vu Thuy Khanh Le-Trilling, Meng-Ji Lu, Dong-Liang Yang, Fei Deng, Ulf Dittmer, Mirko Trilling, Xin Zheng

## Abstract

Approximately half of the SARS-CoV-2 infections occur without apparent symptoms, raising questions regarding long-term humoral immunity in asymptomatic individuals. Plasma levels of immunoglobulin G (IgG) and M (IgM) against the viral spike or nucleoprotein were determined for 25,091 individuals enrolled in a surveillance program in Wuhan, China. We compared 405 asymptomatic individuals with 459 symptomatic COVID-19 patients. The well-defined duration of the SARS-CoV-2 endemic in Wuhan allowed a side-by-side comparison of antibody responses following symptomatic and asymptomatic infections without subsequent antigen re-exposure. IgM responses rapidly declined in both groups. However, both the prevalence and durability of IgG responses and neutralizing capacities correlated positively with symptoms. Regardless of sex, age, and body weight, asymptomatic individuals lost their SARS-CoV-2-specific IgG antibodies more often and rapidly than symptomatic patients. These findings have important implications for immunity and favour immunization programs including individuals after asymptomatic infections.

**One-Sentence Summary:** Prevalence and durability of SARS-CoV-2-specific IgG responses and neutralizing capacities correlate with COVID-19 symptoms.

---

Currently, the world faces a global COVID-19 pandemic. As of March 19, 2021, more than 121.8 million people had a laboratory-confirmed SARS-CoV-2 infection and nearly 2.69 million people died during or in the direct aftermath of COVID-19(*1*). Calculations of the excess mortality and sero-prevalence surveillance programs indicate that the actual numbers of infections and fatalities are far higher. An important determinant for the number of unrecorded cases is the occurrence of very mild and/or asymptomatic infections which are the focus of this study. The scarcity of secondary SARS-CoV-2 infections(*2,3*) indicates that adaptive immune responses prevent re-infections in the vast majority of cases - at least during the approximately one-year period during which SARS-CoV-2 has been studied to date. Others and we have shown that binding and neutralizing antibodies develop rapidly after infection and are maintained in the majority of symptomatic COVID-19 patients for a period of 6-10 months after disease onset(*4–6*). It is important to emphasize that this observation period was defined by the end of the studies rather than the decline of detectable antibodies. However, recent reports suggest that binding antibodies and the neutralizing activity against SARS-CoV-2 is either not similarly strong and/or long-lasting in individuals who had only mild or no symptoms(*7–10*). These important landmark studies either included relatively few patients (e.g., 37 per arm) or only examined a relatively short period of time (e.g., 8 weeks). Additionally, no information concerning possible re-exposures to SARS-CoV-2 was provided. Therefore, we felt that the duration of protective immunity in asymptomatic individuals should be elucidated in larger cohorts and with a more informative study design.

There is a controversial debate concerning the question with which frequencies *bona fide* asymptomatic SARS-CoV-2 infections occur. Another important matter of debate is the question if and how they contribute to the spread of the virus(*11*). A large study from Wuhan suggests that asymptomatic individuals seem not to be very infectious for their contact persons(*12*).

Obviously, unspecific symptoms such as headache, myalgia, and fatigue are not always linked to COVID-19, because they are rather common in the general population and may have various reasons. Thus, the incidence of asymptomatic SARS-CoV-2 infections appears to vary considerably in different studies and/or populations. Descriptions range from 17.8% in Diamond Princess Cruise ship tourists(*13*) to 21.9-35.8% in a nationwide sero-prevalence study in Spain(*14*). Factors influencing this wide range appear to be related to the study design (e.g., retrospective questionnaires), personal expectations of being infected, and maybe the patience and perseverance during interviews and interrogations. Regardless of the actual percentage, two points are beyond doubt: (I) a highly relevant proportion of persons acquires a SARS-CoV-2 infection (as indicated by diagnostic antibody testing) without seeking medical help and without recognizing and/or remembering unusual symptoms, and (II) such asymptomatic individuals have no or far milder symptoms as compared to individuals who actively seek medical help due to the occurrence of symptoms. Thus, asymptomatically SARS-CoV-2-infected individuals are often hard to find for larger immunological studies.

Usually, the timing of asymptomatic infections is uncertain given that the virus itself has never been detected by nucleic acid or antigen testing. In such cases, the retrospective diagnosis is exclusively based on the presence of specific antibodies. Since immunity wanes over time, it is very difficult to accurately determine the prevalence and kinetics of binding and neutralizing antibodies in asymptomatic individuals. In the absence of virus detection and/or symptomatic disease episodes it is nearly impossible to distinguish recent infection events associated with low IgG titers from past infections that had initially elicited strong immune responses which declined afterwards. This level of uncertainty increases even further when re-exposures are taken into account which are almost impossible to detect *in natura* but will almost certainly booster immunity. While the above applies to phases of on-going public virus spread, a clearly defined end of local virus transmission chains may be applied as ‘synchronization element’ since it excludes infections and the associated antigen re-exposure beyond a defined time point. Since April 2020 and despite large-scale public surveillance programs(*15*), no autochthonous virus transmissions have been detected in Wuhan strongly suggesting that the stringent non-pharmacologic interventions virtually terminated local virus spread. Given that this end excludes infections, re-infection, antigen re-exposures, and immunological boostering, we inferred that this serendipitous situation would enable an - at least to our knowledge - unprecedented study design dealing with the aftermath of a COVID-19 endemic. We screened 25,091 outpatients in April 2020 and surveyed antibody responses in more than 987 sero-positive persons during a six-month period after the epidemic in Wuhan had ended. Immunoglobulin M (IgM) and G (IgG) responses recognizing the receptor binding domain (RBD) of the spike (S) or the nucleocapsid (N) protein as well as neutralizing activities of clinical specimens derived from 405 asymptomatically infected individuals and 459 symptomatic COVID-19 patients were determined in a comprehensive and comparative study design. The results provide novel insights into the long-term immune status of asymptomatic individuals and have important implications for the understanding of collective immunity as well as the design of global vaccination programs.

## Methods

### Patients and sample collection

In total, 29,177 clinical specimens obtained from 25,091 outpatients of the clinic of Wuhan Union Hospital during the period between April 2020 and October 2020 were included in this study. The levels of IgM and IgG antibodies recognizing the RBD of the S protein and the N protein (IgG-S, IgG-N, IgM-S, and IgM-N) were determined. A total of 987 individuals who have not been vaccinated against SARS-CoV-2 tested positive for at least one SARS-CoV-2-specific antibody. Focusing on the antibody-positive patients, we conducted interviews to assess whether the persons experienced symptoms such as fever, sore throat, cough, loss of taste or smell, and chest tightness during the epidemic. Of the 987 SARS-CoV-2-specific antibody-positive persons, 123 had to be excluded from further analyses for one or more of the following reasons: refusal to provide medical information, ambiguity of medical information or sole IgM positivity. The latter were excluded because of the limited specificity of IgM responses. Clinical specimens derived from repetitive testing of the same individual during a one-month interval were also not taken into account. Individuals co-infected with human influenza A virus, influenza B virus or other viruses associated with respiratory infections were excluded. In the end, data of 405 asymptomatic persons and 459 symptomatic patients were included in this study (Fig. 1). We retrospectively collected patients’ medical information including demographic factors (Table. S1). Plasma samples were separated by centrifugation at 3000g for 15 min after 30 min-inactivation at 56°C (complement inactivation) and tested concerning the presence of SARS-CoV-2-specific antibodies. All patients signed a general written consent that residual blood samples can be applied for scientific research. All procedures were approved by the Ethics Commission of Union Hospital of Huazhong University of Science and Technology in Wuhan.

**Figure 1:**
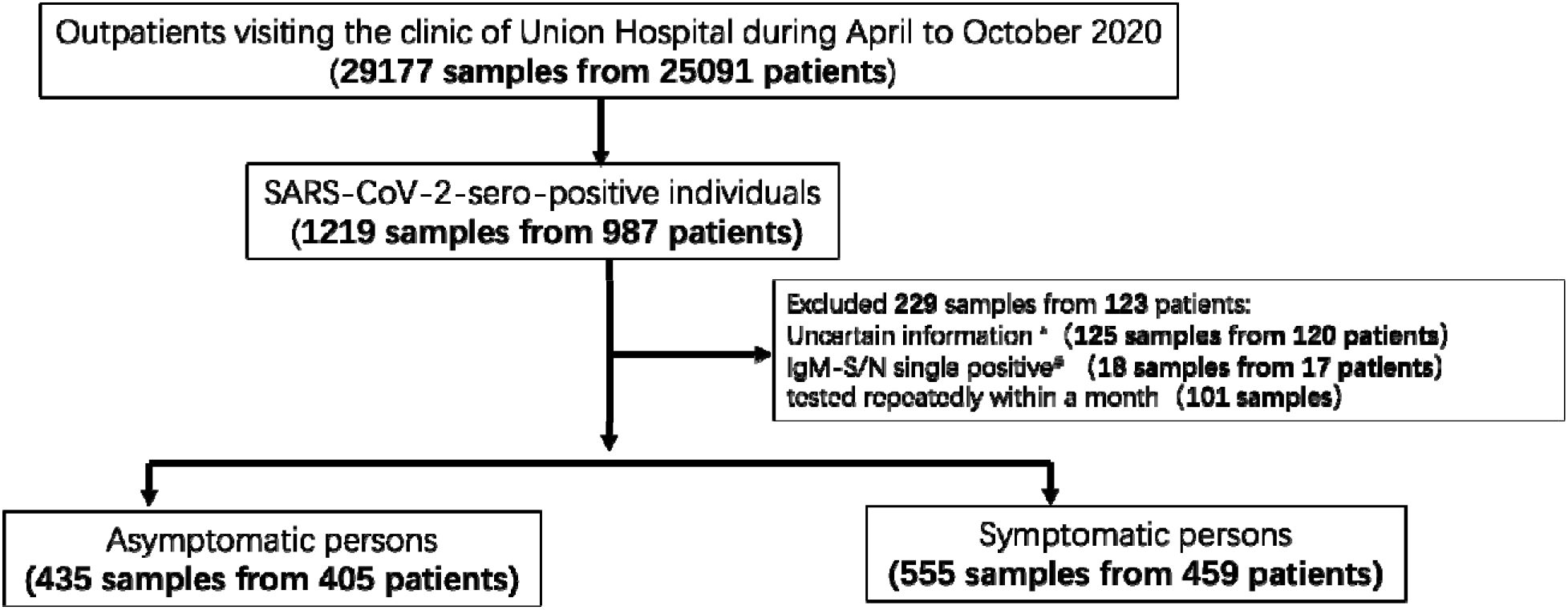
Study population, cohort enrolment process, exclusion criteria, and workflow of surveillance and analyzation. Please note that there was an overlap between exclusion criterion 1* (‘uncertain information’) and 2^#^ (‘IgM single positivity’).

#### Detection of IgG and IgM against spike protein and nucleocapsid protein of SARS-CoV-2

IgG-S, IgG-N, IgM-S, and IgM-N levels were quantified by capture chemi-luminescence immunoassays (CLIA) Kit (Snibe, Shenzhen, China, Lot#: 130219015M/130219016M) using the MAGLUMI™ 4000 Plus as described previously(*6*). The cut-off value for IgM-S was 0.7 AU/mL and 1.0 AU/mL for IgM-N, IgG-S, and IgG-N.

### Virus neutralization test assay

The SARS-CoV-2-neutralizing activity of patient plasma was tested against SARS-CoV-2 (Strain BetaCoV/Wuhan/WIV04/2019, National Virus Resource Center number: IVCAS 6.7512) in highly permissive Vero E6 cells using the described co-incubation methodology(*6*). Virus-specific cytopathic effects (CPE) were visualized and judged by microscopic inspection. The neutralizing antibody titers were expressed as reciprocal value of the highest actual dilution that significantly prevented CPE formation.

### Statistics and reproducibility

The mean and standard deviation were applied for describing continuous variables with a normal distribution. The median and the interquartile range (IQR) were used to describe continuous variables with a skewed distribution. For categorical variables, the number (n) and the percentage (%) were applied for description. We used the Mann-Whitney U test, χ2 test, or Fisher’s exact test as appropriate. A non-parametric Spearman’s correlation test was applied for the correlation analyses. Longitudinal changes in antibody titers during April 2020 and October 2020 were depicted using the locally weighted regression and smoothing scatterplots (Lowess) model (ggplot2 package in R). All reported p values were two-sided, and a p value below 0.05 was regarded as hallmark for statistical significance. Levels of statistical significance were depicted as follows: ns, not significance; *p<0.05; **p<0.01; ***p<0.001; ****p<0.0001. All statistical analyses were conducted using R (The R Foundation, http://www.r-project.org, version 4.0.0).

## Results

The local COVID-19 epidemic in Wuhan was discovered in late 2019(*16*) and lasted until the end of March 2020. During April and May, only seven new cases were identified among more than 11.2 million inhabitants of Wuhan, China. Despite enormous testing efforts (approximately 9.89 million tests were conducted), no autochthonous infections have been identified since June 2020 (Fig. S1). We figured that this temporarily well-defined epidemic may provide an opportunity to determine humoral immune responses elicited by a novel virus infection that necessarily must have occurred during a very precisely defined and narrow timeframe and in absence of subsequent antigen exposures during the post-epidemic period.

Previous virus proteome-wide analyses showed that asymptomatic infections mainly produce IgM and IgG antibodies recognizing the S1 and the N protein of SARS-CoV-2(*8*). Therefore, we focussed on these antibody responses. We determined specific IgG and IgM responses recognizing the N or the S protein, applying capture chemi-luminescence immunoassays (CLIA). More than 29,177 clinical specimens were analysed from 25,091 outpatients who visited the clinic of Union Hospital during the period from April to October 2020. A total of 1,219 plasma specimens obtained from 987 individuals showed at least one type of SARS-CoV-2-specific antibodies, corresponding to an overall sero-prevalence of 3.93% among the participants. This prevalence is highly consistent with two previous studies conducted among Wuhan residents which described sero-positivity rates of 2.39% and 3.9% (*17,18*). After applying reliability criteria such as uncertainty of medical information, repetitive testing during a one-month period, and IgM positivity only (see the M&M section for details), 864 subjects represented by 990 plasma samples were included in this study (Fig. 1 and Table. S1). Interestingly, nearly half of all subjects (n=405; ~46.9%) had no symptoms, whereas 459(~53.1%) suffered from symptomatic COVID-19. The demographic characteristics of those asymptomatic individuals and symptomatic COVID-19 patients were compared. There were no significant differences concerning age, sex or body weight between individuals who experienced symptomatic and asymptomatic infections (Table. S1).

### In the absence of antigen re-exposure, asymptomatic individuals lose SRAS-CoV-2-specifìc IgG-S responses more rapidly than symptomatic patients

As expected, based on the short lifespan of IgM responses and in agreement with the literature(*19*), most plasma-positive individuals, regardless of the presence or absence of symptomatic episodes, did not show IgM responses recognizing SARS-CoV-2-N and S two months after the end of the epidemic (Fig. 2A). Conversely, most individuals initially showed IgG responses recognizing N and S. While both the IgG-N and IgG-S levels remained rather stable during the observation period following symptomatic COVID-19, there was an obvious difference concerning the strength and the sustainability of individual IgG responses compared to asymptomatic individuals, who showed lower and less stable IgG responses (Fig. 2A). Symptomatic patients exhibited an overall IgG-S positivity rate of 89% in April and remained at a prevalence of 80% in October (Fig. 2B). The positivity rate for IgG-N started at 100% in April and remained at 92% during the six-month observation period. The positivity rate for IgG-N in asymptomatically infected individuals also started at 100% and decreased slightly faster to 85% during the observation period (Fig. 2B). However, given the importance of S-specific IgG for protection (e.g., through neutralizing antibodies), we were intrigued by the sharp decline in IgG-S responses (Fig. 2A, lower left panel) and overall positivity rates that dropped from 75% in April to only 49% six months later (Fig. 2B).

**Figure 2:**
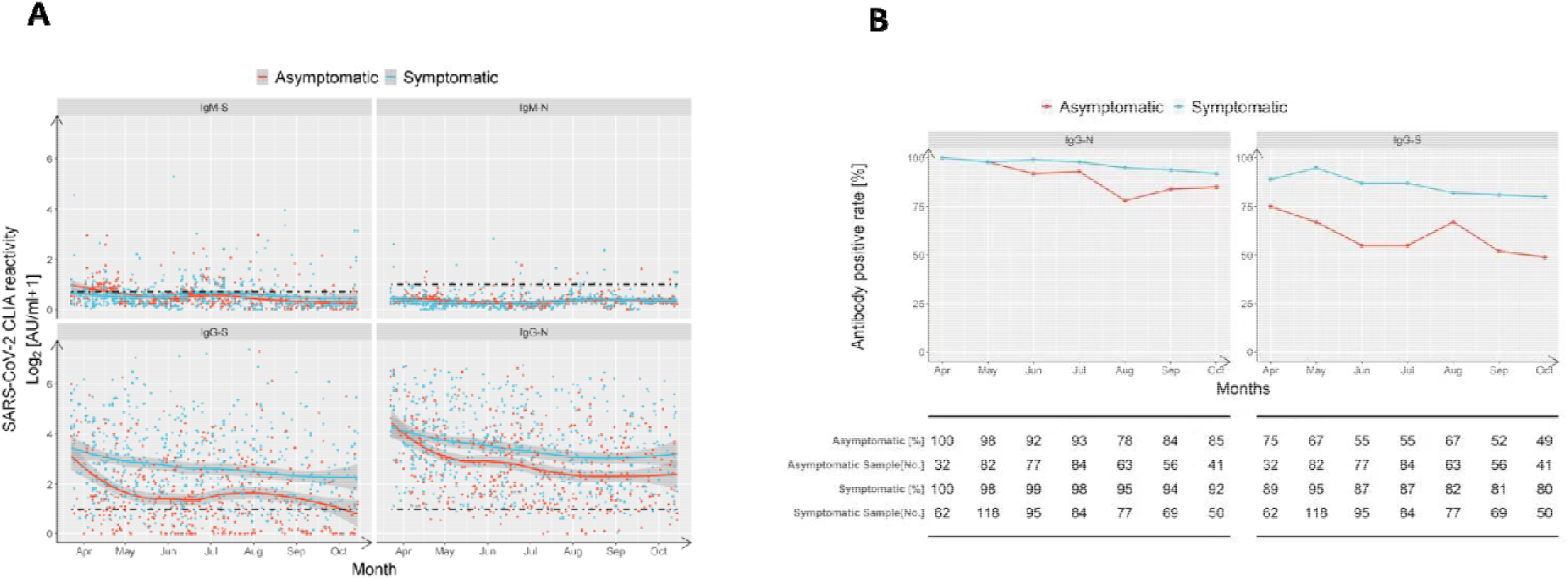
In absence of antigen re-exposure, asymptomatic individuals lose SARS-CoV-2-specific IgG-S responses more rapidly than symptomatic patients. IgM and IgG recognizing the RBD of the spike protein (‘S’) and the nucleoprotein (‘N’) of SARS-CoV-2 were quantified by capture chemi-luminescence immunoassays (CLIA) for 29,177 samples obtained from 25,091 patients. **A**, plasma antibody levels of IgM-S, IgM-N, IgG-S, and IgG-N in asymptomatic (red) and symptomatic (blue) patients obtained during April 2020 and October 2020 are presented. The line shows the mean value calculated using a Lowess regression model and the shaded area represents the 95% confidence interval. **B**, Antibody positivity rates of asymptomatic (red) and symptomatic (blue) groups tested at indicated months after the epidemic ended are shown. The table below the figure depicts the numbers of assessed patients at indicated time points.

### Symptom occurrence is the dominant factor determining the strength and stability of SARS-CoV-2-specific IgG responses

Given these differences in humoral immunity associated with the symptom occurrence, we stratified asymptomatically infected individuals and symptomatic patients according to sex, age, and body mass index (BMI). We then compared the IgG-N and IgG-S antibody levels to examine the influence of symptomatic disease episodes. As shown in Figure 3, IgG-N as well as the IgG-S titers of asymptomatically infected individuals were significantly lower compared to symptomatic patients across most subgroups defined by sex (Fig. 3A), age (Fig. 3B), and BMI (Fig. 3C), except for IgG-N responses in 30-39-year-old and low-weight subjects. We do not think that the lack of significance in the latter two groups indicates a meaningful immunological feature, especially since the trend pointed in the same direction. Taken together, across various groups and biological characteristics of individuals, symptom occurrence during the early phase was the dominant factor defining the strength and sustainability of IgG responses.

**Figure 3:**
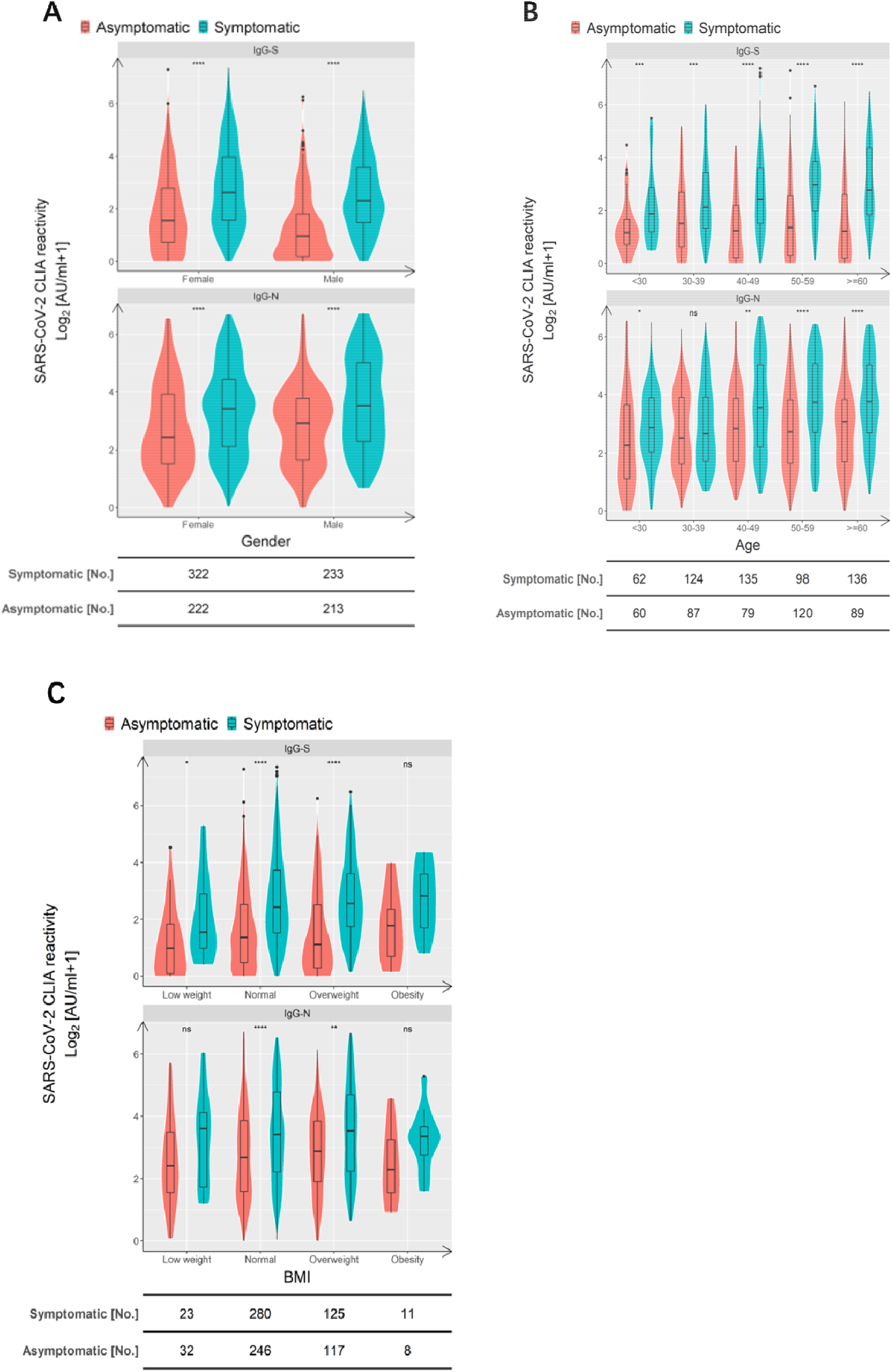
Symptom occurrence is the dominant factor defining the strength of SARS-CoV-2-specific IgG responses. Comparison of RBD S- and N-specific CLIA-reactive IgG titers stratified according to sex (**A**), age (**B**), and BMI (**C**). Violin plots show the distribution of each antibody feature derived from asymptomatic individuals (red) and symptomatic patients (blue). Boxes depict medians (middle line), 75% quartiles (upper bound), and 25% quartiles (lower bound) with whiskers showing a 1.5-fold interquartile ranges above and below boxes. The table below the figure indicates the number of samples obtained from asymptomatic individuals and symptomatic COVID-19 patients. Statistical analysis was performed by a two-tailed Mann-Whitney U test. Asterisks depict the levels of significance as follows: ns, not significant (p>0.05); *p<0.05; **p<0.01; ***p<0.001.

### Neutralization activity is defined by the occurrence of symptoms

Consistent with the essential role of the spike protein for SARS-CoV-2 entry, it represents the main target of neutralizing antibodies. Accordingly, numerous studies including our own documented a strong correlation between IgG-S titers, particularly those antibodies recognizing the receptor binding domain (RBD) of S, and neutralizing activity(*19,20*). The same was observed here: anti-RBD IgG-S titers demonstrated a significant positive correlation with neutralizing activity (Spearman r=0.5795, p<0.0001). A less stringent correlation was found for IgG-N titers (Spearman r=0.1620, p=0.0007) (Fig. S2, left and central panel). Accordingly, high levels of neutralizing activity (1:160 or 1:320) were found in association with high anti-RBD IgG-S levels (Fig. S2, right panel).

Since the IgG-S levels were lower in asymptomatic individuals compared to symptomatic patients, we wondered whether this difference also applies to neutralizing antibodies which are highly relevant for protection from re-infection. We compared neutralizing activities of the two groups at three time points: April, July, and October 2020. As shown in Fig. 4A, the neutralization titers of sera obtained from asymptomatic individuals were significantly lower than those of symptomatic patients. The frequency of individuals showing neutralizing activity in the asymptomatic group showed a downward trend with 59.3%, 51.2%, and 46.3% in April, July, and October, respectively (Fig. 4B). In contrast, the frequency of symptomatic patients with neutralizing activity was stable at a far higher level based on prevalence rates of 77.4%, 86.9%, and 86.0% at the indicated time points (Fig. 4B). From April to July, there was even an increase in the percentage of clinical specimens showing neutralizing antibodies. This may reflect a longterm maturation of antibodies that has been reported after SARS-CoV-2 infection.

**Figure 4:**
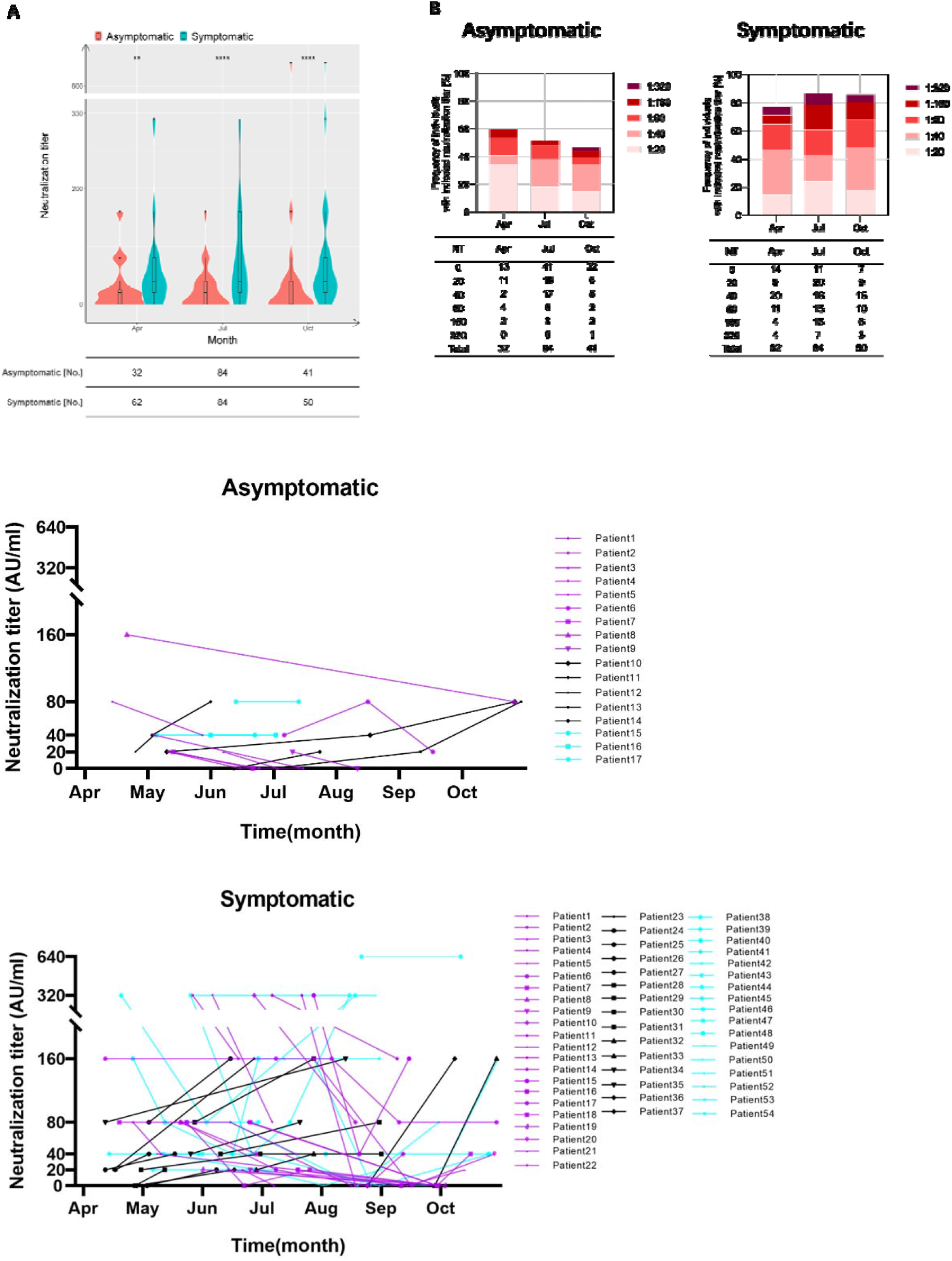
Neutralization activity is defined by the occurrence of symptoms. **A**, Violin plots show the distribution of neutralization titers of sera derived from derived from asymptomatic individuals (red) and symptomatic patients (blue) during April and October 2020. Boxes depict medians (middle line), 75% quartiles (upper bound), and 25% quartiles (lower bound). Whiskers indicate 1.5-times interquartile ranges above and below boxes. The table below the figure highlights the number of clinical specimens that have been assessed. Statistical analysis was performed by two-tailed Mann-Whitney U test. Asterisks depict levels of significance as follows: ns, not significant (p>0.05): *p<0.05; **p<0.01; ***p<0.001. **B**, like in ‘A’ but the frequencies of individuals exhibiting indicated neutralizations titers are depicted. **C**, Sequential sampling and analyses of neutralization activity in 17 asymptomatic individuals and 54 symptomatic COVID-19 patients. Different colours highlight different trends as follows: purple: declining trend; black: increasing trend; light blue: unchanged neutralization.

In order to investigate the neutralizing capacities longitudinally, serum titers of 17 asymptomatically infected individuals and 54 symptomatic patients with repetitive sampling were analysed. Interestingly, the similar proportion of the two groups, 29.4% in symptomatic individuals and 27.7% in symptomatic individuals, showed increasing neutralizing titers over time (Fig. 4C). However, the proportion of individuals with decreasing titers of neutralizing antibodies was 52.9% in asymptomatically infected individuals, but only 40.7% in symptomatic individuals. We found the opposite for individuals with no change neutralization titers over time. Here, the proportion of asymptomatic individuals was only 17.6% in contrast to 31.4 % in symptomatic patients. This indicates that, on the individual level, more patients in the asymptomatic group compared to symptomatic patients show a decrease in their neutralizing capacity over time.

Taken together our data reveal that symptom occurrence during the primary SARS-CoV-2 infection is the dominant factor defining the strength and sustainability of binding and neutralizing IgG antibodies.

## Discussion

We present here, at least to our knowledge, the first comprehensive side-by-side comparison of asymptomatically infected individuals and symptomatic COVID-19 patients in the aftermath of a SARS-CoV-2 endemic. We found striking differences concerning the strength and persistence of SARS-CoV-2-specific IgG responses, in particular in those antibodies recognizing the RBD of S, which comprise neutralizing IgG molecules. Irrespective of sex, age, and body mass index, the symptom occurrence during the early SARS-CoV-2 infection phase was significantly positively correlated with stronger and more sustained IgG-S responses. The same difference was evident concerning the level of neutralizing antibodies.

To enable an investigation such as the present one, several highly unusual circumstances must ‘perfectly’ align: (I) the beginning and the end of the endemic must be well defined, (II) the duration of the endemic needs to be rather short, (III) public surveillance efforts are needed to trace the spread of the virus in the local community, (IV) sufficient numbers of individuals in general and infected subjects in particular need to be present and willing to share their information, and (V) immunologically related viruses must be negligible during the observational period in the studied area. All of these hold true for Wuhan, leading to an unprecedented situation: SARS-CoV-2 was identified here(*16*). The endemic started and ended between late 2019 and end of March 2020 (see Fig. S1). Other seasonal coronaviruses such as HCoV-229E did not circulate extensively during the observational period(*21*) and the antibody responses recognizing the N and S protein of SARS-CoV-2 (if at all) only minimally overlap with responses induced by other seasonal coronaviruses(*22*). In conjunction, these factors allowed us to probe into the strength and sustainability of humoral immunity in the aftermath of a COVID-19 epidemic and in absence of subsequent antigen re-exposure. Importantly, the most decisive factor across different sex, age, and body mass index groups was the occurrence of symptoms during the early SARS-CoV-2 infection.

Albeit in far smaller collectives and in shorter analyses, other authors also observed such difference between patients who experienced severe symptoms and asymptomatic and/or mildly symptomatic groups(*10,23*). We can only speculate why early symptoms correlate with the strength and sustainability of IgG responses. Patient with severe infections may produce higher levels of antibodies during the early disease stage because of the stimulation of a large number of antigens and B cell responses outside germinal centres(*24*). However, damaging effects of SARS-CoV-2 on lymphoid organs affecting the durability of antibody responses and the antibody affinity have also been described(*25,26*). Although several studies reported an association between higher viral loads with more severe symptoms, they found little to no difference in respect to virus loads between pre-symptomatic, asymptomatic, and symptomatic patients(*27*). However, the duration of viral RNA shedding seems to be shorter in people who remain asymptomatic(*11,27*). Thus, a shorter virus replication phase may be associated with an antigen availability that is simply not long enough to prime optimal B cell and/or antibody responses. Another explanation may rely on the association between innate immune responses and symptoms on the one hand and innate immune responses and antibody responses on the other hand. It is well known that interferons, besides their important antiviral activity, by themselves cause flu-like symptoms such as fatigue, fever, and myalgia(*28,29*). Additionally, interferons enhance antibody responses and induce class switching(*30*). Thus, a simple and parsimonious explanation for the association between the occurrence of early symptoms with strong and long-lasting IgG responses may simply be the overlapping dependence on interferons. Since interferon induction is stimulated by viruses, the first explanation (prolonged virus replication) and the second explanation (increased interferon induction) are by no means mutual exclusive.

Some recent studies claimed that neutralizing activities in clinical serum specimens obtained from patients with mild symptoms and/or asymptomatic infections disappears 2 months after infection(*9*). However, other studies(*31*) and our results presented above clearly oppose this view. Despite the apparent decline in antibody responses, nearly half of all asymptomatic individuals exhibited detectable neutralizing activity half a year after the end of the epidemic and in absence of additional antigen exposure. These dynamics are consistent with the change of antibody response during other acute virus infections such as influenza, MERS-CoV, SARS-CoV-1, and the seasonal human coronavirus 229E. Early after such infections, neutralizing antibody titers rise rapidly followed by an obvious contraction phase. However, after this intermediate decline, a stable plateau is established, which can be maintained for several years through the activity of long-lived plasma and memory B cells(*5,26*).

Like all observational analyses, our study has certain limitations. Firstly, after half a year, a potential recall bias of asymptomatic carriers may affect the results of this study. Secondly, a fraction of asymptomatic patients may have been missed by our surveillance as consequence of IgG-S levels below the level of detection during the recovery period(*10,16*). This study only enrolled asymptomatic individuals, who mounted a detectable antibody response.

In conclusion, half a year after the Wuhan COVID-19 epidemic ended, although asymptomatic individuals had lower IgG-S antibody titers, positivity rates, and neutralizing activities compared to symptomatic patients, nearly half of asymptomatic subjects had sufficient neutralization activity. These results suggest that a considerable fraction of asymptomatic natural infections stimulate a humoral immune response conferring the ability to resist reinfections. Despite this good news, above mentioned disparity in the strength and duration of IgG-S responses raised by symptomatic and asymptomatic infections, strongly argue in favour of vaccine programmes including individuals who underwent asymptomatic SARS-CoV-2 infections, ideally with an intermediate prioritization adjusted between vulnerable uninfected individuals and symptomatic COVID-19 patients.

## Supplementary Figures

**Suppl. Figure 1:**
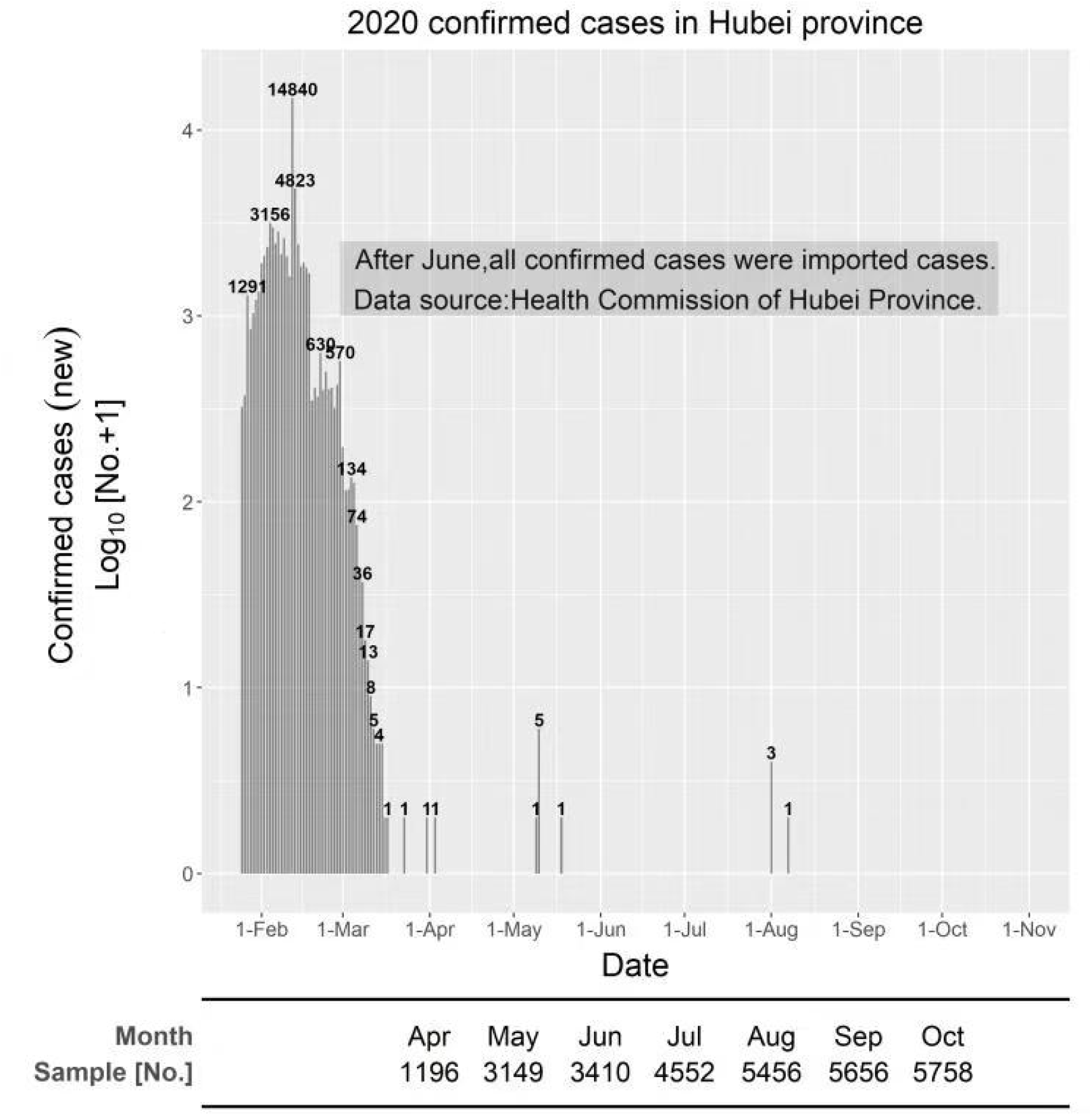
The local spread and the COVID-19 endemic ended before April 1^st^ 2020. The graph depicts newly confirmed SARS-CoV-2 infections in the Hubei province comprising Wuhan according to the public information released by the Health Commission of Hubei Provincial. The table shows the number of samples tested every month.

**Suppl. Figure 2:**
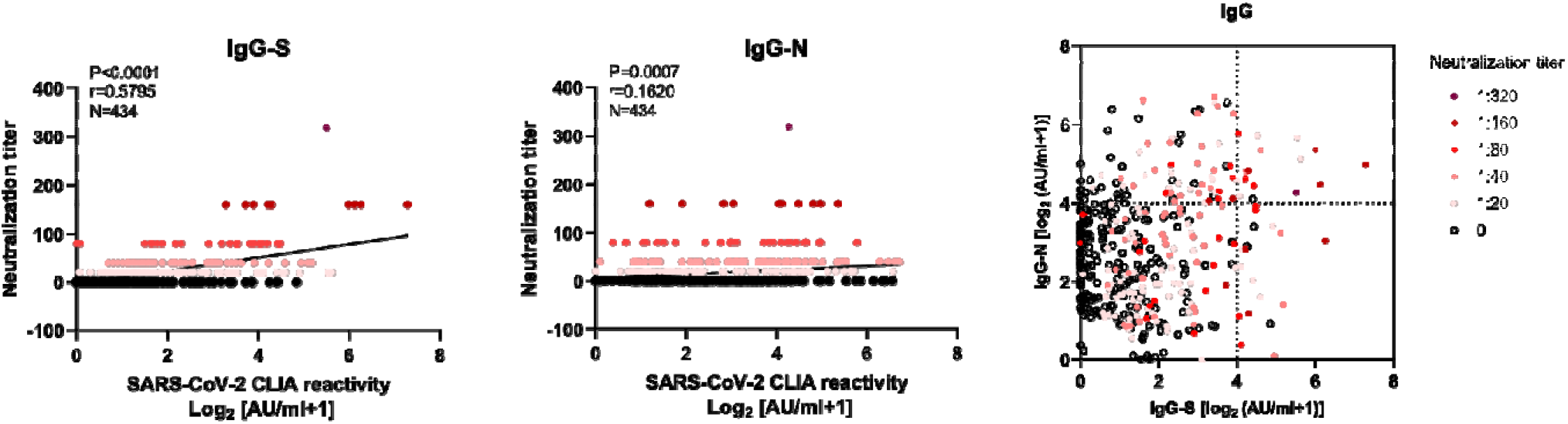
Neutralizing antibodies correlate best with IgG recognizing the RBD of the SARS-CoV-2 spike protein. **A**, Correlation analysis of neutralization titers versus S-specific (left panel) and N-specific (right panel) CLIA-reactive IgG in asymptomatic individuals. A non-parametric Spearman’s correlation test was applied for statistical analyses. In the graphs, p, r, and N indicate the p value, the correlation coefficient, and the sample size, respectively. **B**, Distribution of neutralizing activity as indicated by the colour at different levels of IgG-S (x axis) and IgG-N (y axis).

**Suppl. Table 1.** Demographic characteristics of asymptomatic individuals and symptomatic COVID-19 patients. All data are presented as median with the interquartile range or as number and percentage. Statistic testing was conducted applying a χ2 test (marked by *) or a Mann-Whitney U test (marked by #). A p value <0.05 was considered as hallmark of statistical significance.

|  |  | Total (N=864) | Asymptomatic (N=405) | Symptomatic (N=459) | P-value |
| --- | --- | --- | --- | --- | --- |
| Gender | Male | 387 (44.79%) | 195 (48.15%) | 192 (41.83%) | 0.5052 * |
|  | Female | 477 (55.21%) | 210 (51.85%) | 267 (58.17%) | 0.5468 * |
| Age | Mean (SD) | 47 (14) | 47 (14) | 47 (14) | 0.9027 # |
|  | <30 | 106 (12.27%) | 54 (13.33%) | 52 (11.33%) | 0.6871 * |
|  | 30-39 | 185 (21.41%) | 79 (19.51%) | 106 (23.09%) | 0.5833 * |
|  | 40-49 | 178 (20.60%) | 72 (17.78%) | 106 (23.09%) | 0.4062 * |
|  | 50-59 | 197 (22.80%) | 114 (28.15%) | 83 (18.08%) | 0.1386 * |
|  | >=60 | 198 (22.92%) | 86 (21.23%) | 112 (24.40%) | 0.6389 * |
| BMI | Low weight | 52 (7.04%) | 32 (8.56%) | 20 (5.48%) | 0.4111 * |
|  | Normal | 464 (62.79%) | 228 (60.96%) | 236 (64.66%) | 0.7413 * |
|  | Overweight | 205 (27.74%) | 107 (28.61%) | 98 (26.85%) | 0.8132 * |
|  | Obesity | 18 (2.44%) | 7 (1.87%) | 11 (3.01%) | 0.6058 * |

\* Calculated using the $\chi^2$ test.

## Funding

National Science and Technology Major Project for Infectious Diseases of China 2018ZX10302206 (XZ)

National Science and Technology Major Project for Infectious Diseases of China 2018ZX10723203 (XZ)

the Applied Basic and Frontier Technology Research Project of Wuhan 2020020601012233 (XZ)

the Fundamental Research Funds for the Central Universities 2020kfyXGYJ016 (XZ)

the Fundamental Research Funds for the Central Universities 2020kfyXGYJ028 (JL)

the Tongji-Rongcheng Center for Biomedicine, Huazhong University of Science and Technology, the Medical Faculty of the University of Duisburg-Essen, and Stiftung Universitätsmedizin Essen, University Hospital Essen, Germany (DY)

the Kulturstiftung Essen and the Deutsche Forschungsgemeinschaft (DFG) through grants RTG 1949/2 (MT)

the Kulturstiftung Essen and the Deutsche Forschungsgemeinschaft (DFG) through grants TR1208/1-1 (MT)

the Kulturstiftung Essen and the Deutsche Forschungsgemeinschaft (DFG) through grants TR1208/2-1 (MT)

## Author contributions

Each author’s contribution(s) to the paper should be listed [we encourage you to follow the <u>CRediT</u> model]. Each CRediT role should have its own line, and there should not be any punctuation in the initials.

Conceptualization: JW, BYL, JL, FD, UD, MT, XZ
Methodology: BYL, YHF, HW, XLY, SS, LKC, SML, SHL, TDX,
Investigation: YHF, HW, XLY, SS, SML, SHL, TDX,
Visualization: JW, BYL, HW, XLY, LKC, VTKLT,
Funding acquisition: JL, DLY, MT, XZ
Project administration: SS, JL, MJL, DLY, FD, UD,
Supervision: JW, MJL, DLY, UD, MT, XZ
Writing – original draft: JW, BYL, YHF, HW, XLY, VTKLT, MT, XZ
Writing – review & editing: JW, MJL, FD, UD, MT, XZ

## Data and materials availability

We can provide participant data without names and identifiers for the purpose of protecting the patients’ privacy. During the review period of the article, we can provide all the original data for reviewers. After the article is accepted, we will upload the data to the public database.

